# *KARRIKIN INSENSITIVE2* (*KAI2*)-dependent signaling pathway controls vegetative reproduction in *Marchantia polymorpha*

**DOI:** 10.1101/2022.11.29.518368

**Authors:** Aino Komatsu, Kyoichi Kodama, Yohei Mizuno, Mizuki Fujibayashi, Satoshi Naramoto, Junko Kyozuka

## Abstract

In vegetative reproduction of *Marchantia polymorpha*, propagules, called gemmae, are formed in gemma cups. Despite its significance for survival, control of gemma and gemma cup formation by environmental cues is not well understood. We show here that the number of gemmae formed in a gemma cup is a genetic trait. Gemma formation starts from the central region of the floor of the gemma cup, proceeds to the periphery, and terminates when the appropriate number of gemmae are initiated. The MpKARRIKIN INSENSITIVE2 (MpKAI2)-dependent signaling pathway promotes gemma cup formation and gemma initiation. The number of gemmae in a cup is controlled by modulating the ON/OFF switch of the KAI2-dependent signaling. Termination of the signaling results in the accumulation of MpSMXL, a suppressor protein. In the Mp*smxl* mutants, gemma initiation continues, leading to the formation of a highly increased number of gemmae in a cup. Consistent with its function, the MpKAI2-dependent signaling pathway is active in gemma cups where gemmae initiate, as well as in the notch region of the mature gemma and midrib of the ventral side of the thallus. In this work, we also show that *GEMMA CUP-ASSOCIATED MYB1* works downstream of this signaling pathway to promote gemma cup formation and gemma initiation. We also found that the availability of potassium affects gemma cup formation independently from the KAI2-dependent signaling pathway in *M. polymorpha*. We propose that the ancestral function of the KAI2-dependent signaling pathway may be to optimize vegetative reproduction by adapting to the environment.

## Introduction

Production of new individuals from somatic cells is generally called vegetative or asexual reproduction [1]. Many plants demonstrate vegetative reproduction as the prevailing way of propagation, such as reproduction through bulbs, tubers, and rhizome buds. Plantlets can also initiate from leaf margins in some species, such as *Kalanchoe daigremontiana*.

Bryophytes propagate through both sexual and vegetative reproduction [2, 3]. In the vegetative reproduction of *Marchantia polymorpha*, propagules called gemmae are formed in a cup called a gemma cup. Gemma cups derive from the apical stem cell and are regularly developed in conjunction with the bifurcation of the apical notch [4, 5]. The gemmae are dormant in the gemma cup and start to grow after they leave the cup. Hundreds of gemmae are formed in a gemma cup, and each gemma has the potential to grow as an individual, making vegetative reproduction in *M. polymorpha* a highly efficient and vigorous means of propagation. In general, the level of vegetative propagation needs to be appropriately regulated to keep a balance between propagation and resources available. However, the regulation of vegetative reproduction and its underlying mechanisms in *M. polymorpha* is not well understood.

Recently, regulators of vegetative propagation of *M. polymorpha* have been identified. *GEMMA CUP-ASSOCIATED MYB1* (*GCAM1*) encoding an MYB transcription factor is essential for gemma cup initiation [6]. The orthologs of *GCAM1* in *Arabidopsis* and tomato were shown to be regulators of axillary meristem formation [7–10]. This suggests that basic mechanisms are shared between gemma cup formation in *M. polymorpha* and axillary meristem formation in angiosperms. Mp*CLE2*, the closest homolog of *CLAVATA3* (*CLV3*) of *Arabidopsis*, affects gemma cup formation through a conserved CLV3-CLV1-CIK module [11, 12].

Gemmae initiate from the bottom of the gemma cup [2, 5]. Some of the gemma cup floor cells are specified as gemma initial cells or mucilage cells. The cells specified as the gemma initial cells continue divisions and become gemmae. The mechanisms underlying the cell fate specification remain to be understood. In the mutants of *KARAPPO* (*KAR*), encoding an ROP guanine nucleotide exchange factor (Rop GEF), gemma initial cells are not specified, while gemma cups and mucilage cells normally differentiate [13]. This indicates that KAR is essential to specify cells as gemma initial cells, and distinct mechanisms control gemma initiation and gemma cup formation. This is also supported by the fact that no defects in gemma initiation are observed in the Mp*cle2* mutants [11].

Plant hormones are involved in the control of gemma cup and gemmae formation. Reduction of cytokinin levels by over-expression of the cytokinin oxidase gene Mp*CKX2* completely suppressed gemma cup formation, indicating that cytokinins are essential for gemma cup formation [14, 15]. However, it is not known whether cytokinin is also required for gemma initiation because the gemma cup was absent in the CKX over-expression lines as in *gcam1*, in which gemma cup formation is wholly blocked. Auxin likely plays an opposite role to the cytokinins in the control of gemma cup formation. Although detailed mechanisms remain to be elucidated, genetic analysis and pharmacological experiments indicated that auxin plays a negative role in gemma cup formation through an AUX/IAA-ARF-dependent signaling pathway [16, 17].

*KARRIKIN INSENSITIVE 2* (*KAI2*) is an ancient ortholog of *DWARF14* (*D14*), the receptor of strigolactones (SLs), a class of plant hormones [18–21]. In the SL signaling pathway, D14 interacts with MORE AXILLARY GROWTH2 (MAX2)/D3, an F-box protein, upon ligand binding [21, 22]. This leads to interaction with D53/SMXL6,7,8, suppressor proteins, resulting in their degradation [23–26]. KAI2 was identified as a signaling component of karrikins (KARs), butenolides derived from burned plants, and induce seed germination [19, 27–29]. Later, it was shown that KAI2 also perceives unidentified endogenous ligand(s) tentatively called KAI2 ligands (KL) [19, 30]. The KAR/KL signal is transduced through a pathway similar to the SL signaling pathway. MAX2 is shared between the two pathways, and MAX2-dependent proteasome-mediated degradation of suppressor proteins is crucial to controlling downstream genes in both pathways. In *Arabidopsis*, SUPPRESSOR OF MAX1 (SMAX1) and SMAX1 LIKE2 (SMXL2) are suppressor proteins degraded by MAX2 upon ligand perception by KAI2 [28, 31, 32]. SMAX1, SMXLs, and D53 belong to the same family of proteins with weak homology to ClpB-type chaperonins [23, 24, 33]. The KAR/KL signaling pathway in seed plants controls diverse aspects of growth and development, including seed germination, seedling growth, leaf development, root growth, root hair development, stress tolerance, and arbuscular mycorrhizal symbiosis [18, 28, 31, 34–39]. Generation of D14 by duplication of KAI2 occurred in the common ancestor of seed plants; thus, D14 is absent in the bryophytes [19, 33, 40, 41]. *M. polymorpha* contains two *KAI2* paralogues (Mp*KAI2A* and Mp*KAI2B*), one *MAX2* gene (Mp*MAX2*), and one *SMXL* (Mp*SMXL*) [42]. We previously reported that MpKAI2A, MpMAX2, and MpSMXL work in a single signaling pathway, which depends on the degradation of MpSMXL [43]. KAI2-dependent signaling in *M. polymorpha* is involved in the control of thallus growth and gemma dormancy [43].

In this work, we show that the number of gemmae in a gemma cup is a genetically controlled trait. The MpKAI2-dependent signaling pathway positively controls gemma cup formation and gemma initiation. It is turned off when the appropriate number of gemmae has been initiated in the gemma cup. Thus, the number of gemmae in a cup is controlled by modulating the ON/OFF switch of the KAI2-dependent signaling.

## Results

### KAI2-dependent signaling promotes gemma cup formation

Gemma cups are regularly produced on the midrib region of the dorsal side of the thallus (Figure 1A). Gemma cups and midrib derive from the dorsal merophyte generated from the apical cell [2, 5]. The first signs of gemma cup differentiation can be observed a few cells away from the apical cell. The location of the developing gemma cup moves towards the proximal direction along the thallus growth. The apical region of each thallus bifurcates due to duplication of the apical cell, resulting in duplication of the potential gemma formation points. The pattern of apical cell duplication, thallus bifurcation, and gemma cup formation is constant in our culture conditions, as shown in Figure 1A. Typically, one gemma cup is produced from the apical cell until the next bifurcation. Previously, we reported that mutations in Mp*kai2a* and Mp*max2*, the KAI2-signaling components caused retarded and upward growth of thalli and reduced dormancy of gemma under dark conditions [43]. In addition to these defects, we observed that gemma cup formation was strongly suppressed in Mp*kai2a* and Mp*max2* loss of function mutants grown on soil (vermiculite) (Figures 1B, 1C and S1). On the other hand, these mutants normally formed gemma cups when grown on half-strength Gamborg’s B5 medium (1/2 B5 medium). Gemma cup formation was not significantly affected in the Mp*smxl1*, regardless of medium or soil.

**Figure 1.**
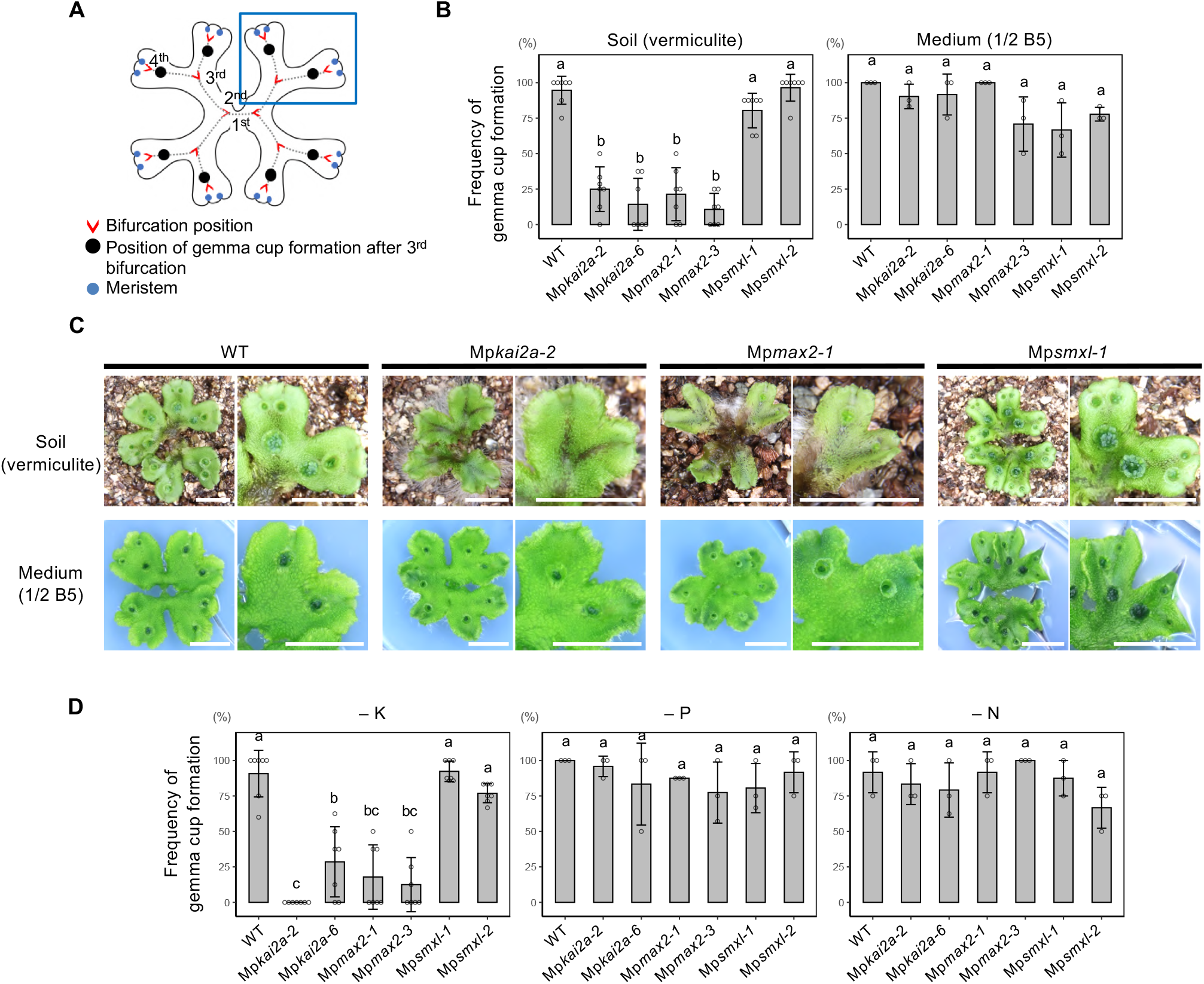
KAI2-dependent signaling promotes gemma cup formation. (A). Schematic diagram of a *M. polymorpha* plant, and the position of the gemma cups on the thallus. The numbering indicates the order of thallus bifurcation. Gemma cups formed after the third bifurcation (black dots) were analyzed in (B) and (D). See also Figure S1. (B). Frequency of actual gemma cup formation per eight possible gemma cup formation points after 3^rd^ bifurcation in plants grown on soil (left) and synthetic medium (right). Bars represent mean ± SD (n = 7 for soil, n = 3 for synthetic medium). (C). Gemma cup phenotypes of KAI2-dependent signaling mutants grown on soil or synthetic medium. Plants were grown from a gemma on medium (1/2 B5) for six days, then transferred to soil (upper panel) or a fresh 1/2 B5 medium (lower panel) and grown for another 14 days. Scale bars; 1 cm. (D). Frequency of actual gemma cup formation per eight possible gemma cup formation points after 3^rd^ bifurcation grown on 1/2 B5 medium lacking potassium (-K), phosphate (-P), or nitrogen (-N). Plants were grown from a gemma on 1/2 B5 medium for six days, then transferred to the respective nutrient-deficient medium and grown for another 14 days. Bars represent mean ± SD (n = 7 for (-K) and (-P), n = 3 for (-N)). See also Figure S2. The Tukey’s HSD test was used for multiple comparisons in (B) and (D). Statistical differences (p < 0.05) are indicated by different letters.

Since gemma cup formation is more severely affected in plants grown on the soil than in those on the medium, we assumed that gemma cup formation is affected by nutrient availability. To test this hypothesis, we grew the KAI2 signaling mutants on 1/2 B5 medium lacking potassium (K), phosphate (P), or nitrate (N) (Figures 1D, S2 and S3). Depletion of these nutrients did not abolish gemma cup formation in WT. However, gemma cup formation was strongly suppressed in the Mp*kai2a* and Mp*max2* mutants by the lack of K. Although the overall growth of plants was retarded, gemma cup formation was not affected by the shortage of P or N in Mp*kai2a* and Mp*max2*. Gemma cup formation was suppressed when both the K pathway and KAI2-dependent signaling were blocked. We previously showed that *DIENLACTONE HYDROLASE PROTEIN1* (*DLP1*), encoding an αβ-hydrolase protein, and Mp*KAI2B* are sensitive markers of the KAI2-dependent signaling activity [43]. Expression of *DLP1* and Mp*KAI2B* was unchanged in limiting K conditions (Figure S4), supporting the notion that KAI2-dependent signaling and K independently and redundantly control gemma cup formation.

### KAI2A-dependent signaling pathway positively controls gemmae initiation

We tested the possibility that the KAI2-dependent signaling is involved in the control of gemma formation. As a first step, we asked if the number of gemmae in a gemma cup is genetically defined. We found that a relatively constant number of gemmae are formed in a gemma cup in WT plants in a given growth condition, suggesting that the number of gemmae formed in a gemma cup is a genetically defined trait. We showed that gemma cup formation was not affected in Mp*kai2a* and Mp*max2* when plants were grown on 1/2 B5 medium (Figure 1B), whereas the number of gemmae in a gemma cup was reduced in Mp*kai2a* and Mp*max2* (Figure 2A, B). In contrast, the number of gemmae in a gemma cup was increased in Mp*smxl* loss-of-function mutants (Figure 2A, B). MpSMXL works as a repressor of the downstream responses in KAI2 signaling in *M. polymorpha* [43]. Thus, in the absence of the MpSMXL function, suppression of downstream genes is canceled, mimicking the maximal signaling state. The increase in the gemma number in the Mp*smxl* indicates that the KAI2-dependent signaling promotes gemma initiation through the degradation of MpSMXL, which suppresses the factors that promote gemma formation.

**Figure 2.**
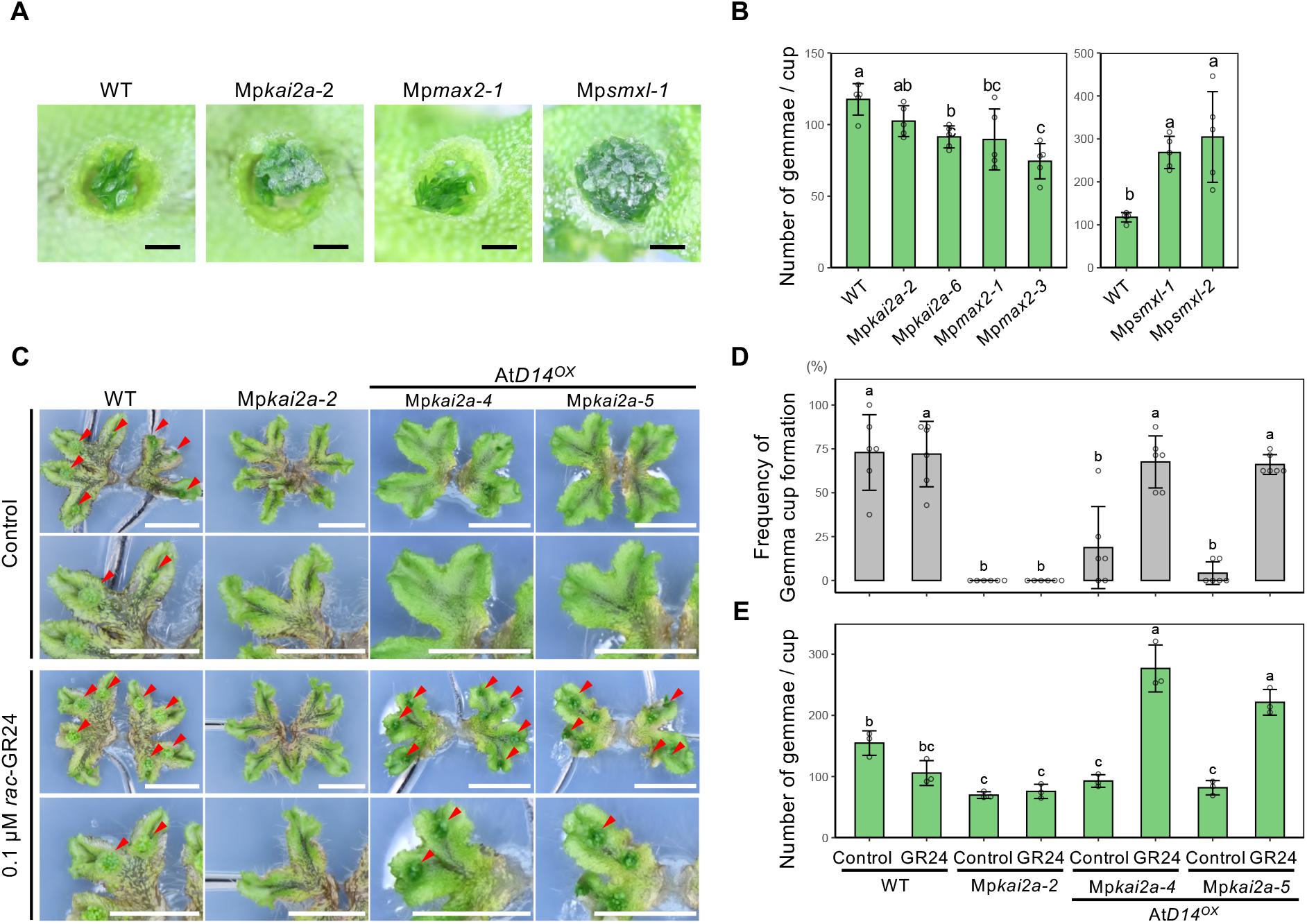
KAI2A-dependent signaling pathway positively controls gemma initiation. (A). Top view of Day 14 gemma cups of WT, Mp*kai2a-2*, Mp*max2-2*, and Mp*smx-1*. (B). The number of gemmae in Day 14 gemma cups of WT, Mp*kai2a-2*, Mp*max2-2*, and Mp*smx-1*. Bars represent mean ±SD from five gemma cups. (C), (D) and (E). Effect of *rac*-GR24 application on morphology of plants (C), frequency of gemma cup formation (D), and the number of gemmae per cup (E). Gemmae of WT, Mp*kai2a-2*, and At*D14^OX^*/ Mp*kai2a-2* were grown on 1/2 B5 medium for six days, then transferred to a potassium-deficient medium and grown for another 21 days with the application of *rac*-GR24 (0.1 μM) every other day. Red arrowheads in (C) indicate gemma cups. Bars represent mean ± SD (n = 6 in (D) and n = 3 in (E). The Tukey’s HSD test was used for multiple comparisons in (B), (D), and (E). Statistical differences (p < 0.05) are indicated by different letters. Scale bars: 1 mm (A), 1 cm (C).

We previously reported that the introduction of At*D14* gene, the strigolactone receptor of *Arabidopsis*, confers responsiveness to SLs through signaling with MpMAX2 and MpSMXL in *M. polymorpha* [44]. Because *M. polymorpha* does not synthesize SLs, At*D14*^ox^ lines can be used as a sensitive inducible system of the KAI2-dependent signaling pathway by adding *rac*-GR24, a synthetic analog of the SL. We used the At*D14*^ox^ lines to confirm that gemma cup formation and gemma initiation are under the control of the MpKAI2A-dependent signaling pathway. When plants were grown on 1/2 B5 medium lacking K, gemma cup formation was severely suppressed in Mp*kai2a* and At*D14^ox^* Mp*kai2a* lines (Figures 2C, D). In addition, the number of gemmae in a gemma cup was also reduced in these lines (Figures 2C, E). These defects were rescued in At*D14^ox^* Mp*kai2a* lines but not in Mp*kai2a* by applying *rac*-GR24, indicating that the control of gemma number and gemma formation is under the control of the KAI2-dependent signaling pathway.

### KAI2-dependent signaling is active in the gemma cup, gemmae, and the mid-rib

We examined the spatial localization of the KAI2-dependent signaling activity. KL, the ligand of the signaling pathway, remains unknown; however, it is anticipated that MpKAI2A works as its receptor in this signaling pathway [43]. Mp*KAI2B*, encoding an αβ-hydrolase protein homologous to MpKAI2A, is strongly up-regulated by KAI2 signaling. Thus, it is a suitable marker of the signaling activity [43]. In order to clarify the spatial localization of the KAI2-dependent signaling activity, we produced transgenic plants containing the *Citrine* gene driven by either the Mp*KAI2A* promoter (_pro_*KAI2A-Cit*) or the Mp*KAI2B* promoter (_pro_*KAI2B-Cit*). Among around twenty independent lines containing each marker gene, two representative lines for each construct were chosen for further analysis. On the dorsal side of the thallus, strong Citrine fluorescence was detected in gemma cups in both _pro_*KAI2A-Cit* and _pro_*KAI2B-Cit* lines under a fluorescence microscope (Figures 3A to C). In addition to the signal in gemma cups, a weaker signal was observed on the entire dorsal surface of the thallus. Detailed observations in transverse sections showed the localization of the fluorescence in the wall and the floor cells of the gemma cup and initiating gemmae in both lines (Figures 3D, E). On the ventral side of the thallus, the Citrine fluorescence was observed in the epidermal cells in the midrib region and ventral scales in both _pro_*KAI2A-Cit* and _pro_*KAI2B-Cit* lines (Figures 3D, E).

**Figure 3.**
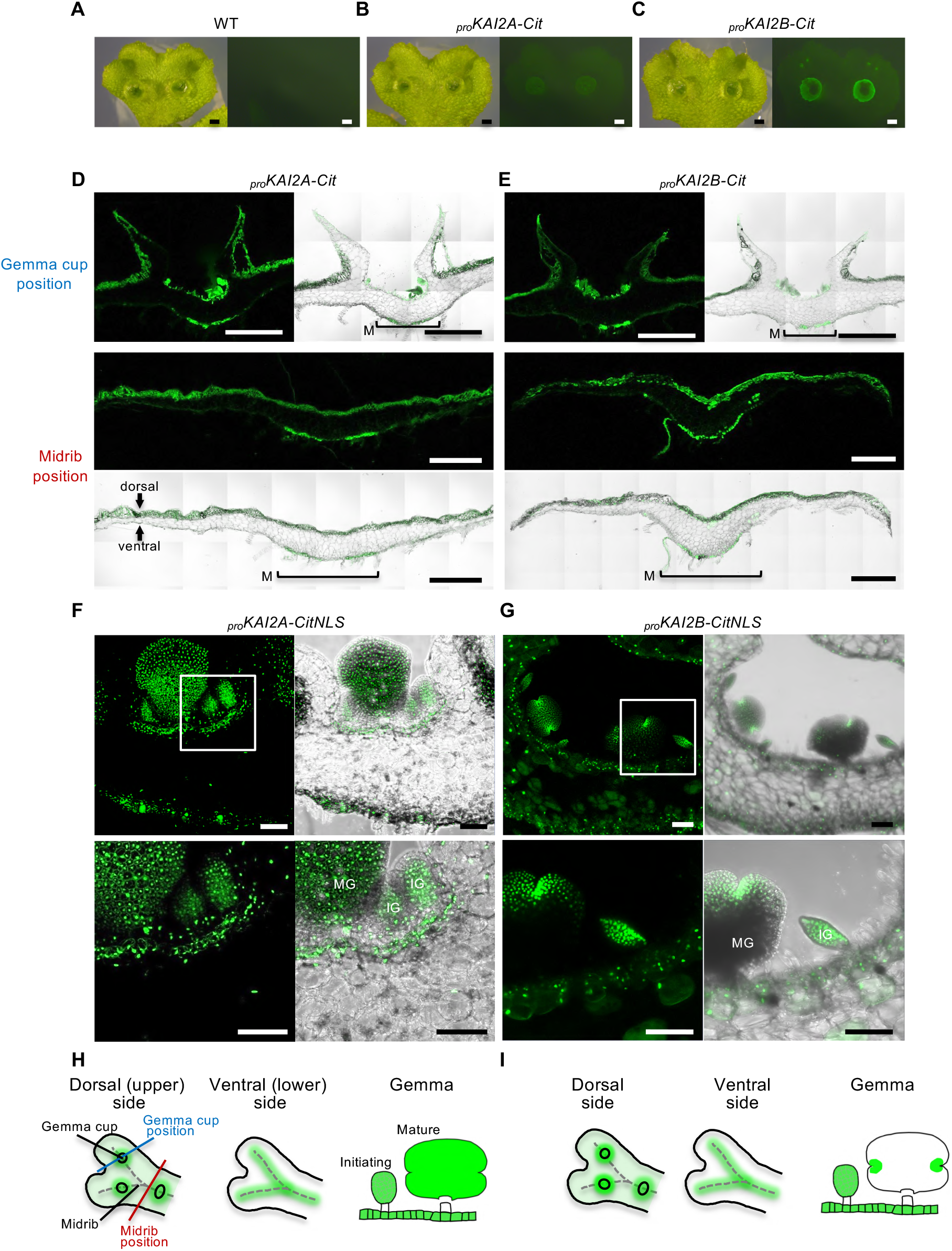
KAI2-dependent signaling is active in the gemma cup, gemmae, and the mid-rib. (A)-(C). Localization of citrine fluorescence in the thallus of WT (A), *_pro_*Mp*KAI2A-Cit* plants (B), and *_pro_*Mp*KAI2B-Cit* plants (C), observed by fluorescence stereomicroscope. (D), (E). Localization of citrine fluorescence in the cross-sections of the gemma cup- (top) and midrib- (bottom) of *_pro_*Mp*KAI2A-Cit* (D) and *_pro_*Mp*KAI2B-Cit* (E) plants observed by confocal laser-scanning microscopy. M: midrib region (F), (G). Localization of citrine fluorescence in the cross-sections of the bottom of the gemma cups and initiating gemmae in the *_pro_*Mp*KAI2A-Cit* (F) and *_pro_*Mp*KAI2B-Cit* (G) lines (top). Enlarged views of the framed regions are shown (bottom). IG: immature gemma, MG: mature gemma (H), (I). Schematic diagram of the Mp*KAI2A* (H) and Mp*KAI2B* expression (I). Scale bars: 1mm (A to E), 100 μm (F, G).

We further examined the spatial localization of the signaling activity using lines expressing the *Citrine* fused with a nuclear localization signal (NLS) driven by the promoter of Mp*KAI2A* (_pro_*KAI2A-CitNLS*) or Mp*KAI2B* (_pro_*KAI2B-CitNLS*). Analysis using these lines showed that the fluorescence was restricted to one or two layers of the gemma cup floor cells and the ventral midrib (Figures 3F, G). During gemma initiation, the fluorescence was observed in the whole region around the initiating gemmae in both _pro_*KAI2A-CitNLS* and _pro_*KAI2B-CitNLS* lines, and this pattern was maintained in _pro_*KAI2A-CitNLS* (Figure 3F). On the other hand, the fluorescence became confined to the notch region as the gemma grows in _pro_*KAI2B-CitNLS* lines (Figure 3G). In conclusion, the localization of the MpKAI2A receptor mostly overlaps with that of MpKAI2B, the downstream marker gene. Both localize in the whole region of the gemma cup, the epidermis of the dorsal side of the thallus, the midrib of the ventral side, and the initiating gemmae, indicating that the KAI2-dependent signaling is active in these regions (Figure 3H, I). A clear difference is that the receptor is expressed in the whole region of the gemmae throughout their formation, while the activity of the signaling pathway becomes restricted to cells around the apical notch as the gemma grows.

### Gemma initiation terminates when an appropriate number of gemmae is reached in a gemma cup

We sought to understand how the increased number of gemmae are formed in the Mp*smxl* mutant. We assumed that a cause for the increase in the number of gemma may be accelerated or prolonged gemma initiation. To clarify this, we first analyzed the time course of gemma accumulation in a gemma cup in WT and Mp*smxl* mutants. In this analysis, the first day when the gemma cup became visible to the naked eye, at the tip of the thallus, was set as Day 1 and the gemma number was counted over time. The pattern of gemma accumulation was similar between WT and Mp*smxl* mutants for the first several days (Figure 4A). In WT, the number of gemmae increased for the first several days and leveled off at around Day 10 and stopped after Day 14, while it continued to increase even after 35 days in Mp*smxl*. This indicates that MpSMXL is required to prevent gemma initiation over time. Since MpSMXL is degraded by the activation of the KAI2 signaling pathway [43], we hypothesized that the KAI2-dependent signaling is active for the first ten days to promote gemma initiation and stops when a certain number of gemma is initiated in a cup in WT. This would lead to the accumulation of MpSMXL, resulting in the prevention of further gemma initiation.

**Figure 4.**
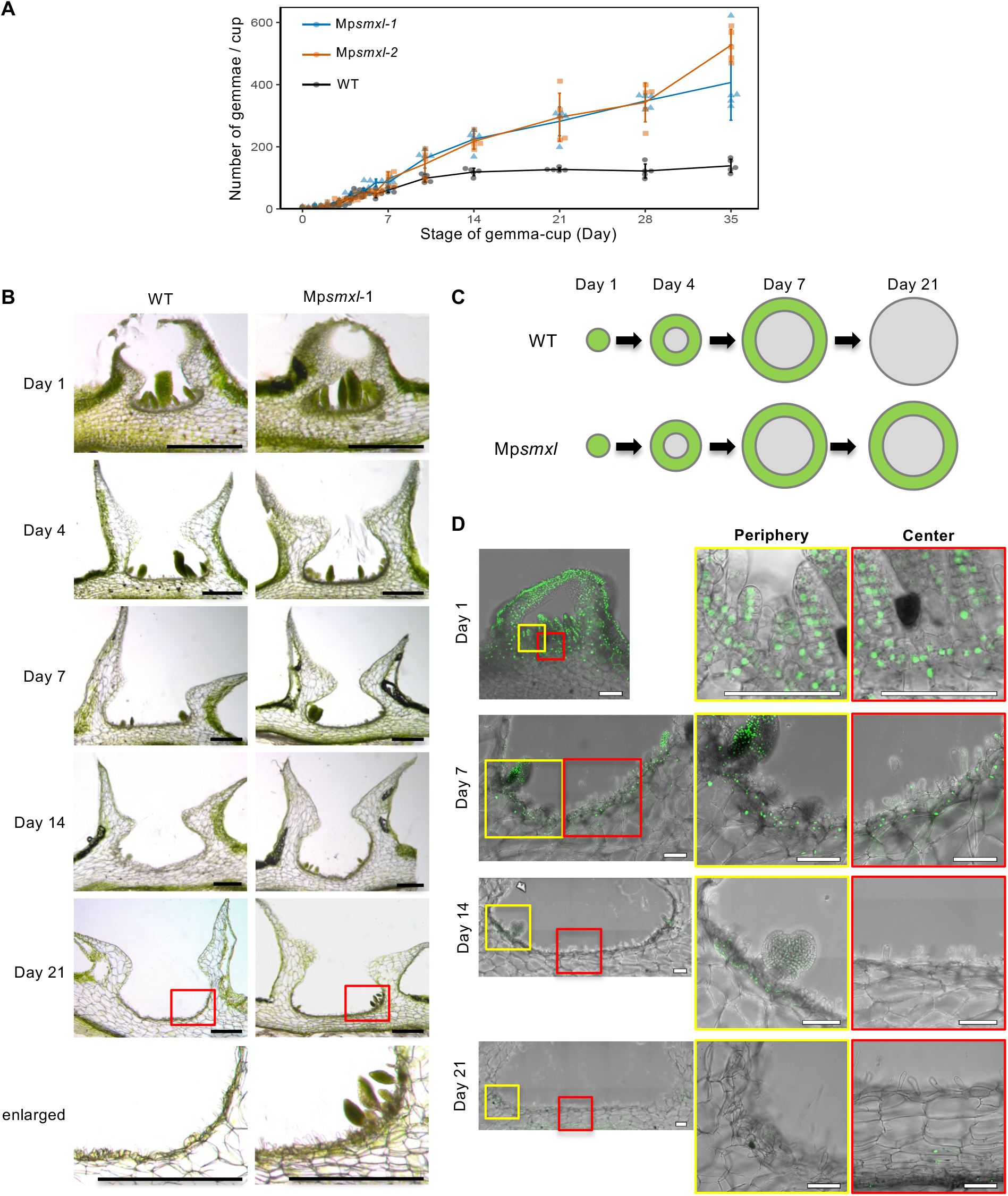
Gemma initiation terminates when an appropriate number of gemmae is reached in a gemma cup. (A). Time-course of gemma accumulation in gemma-cups of WT (black), Mp*smxl-1* (blue), and Mp*smxl-2* (red). The first day when the gemma cup becomes visible is designated as Day 1. Means with the ±SD (n = 5) are indicated. (B). Cross-sections of the gemma cups on Days 1, 4, 7, 14, or 21 of WT and Mp*smxl-1*. Enlarged views of the peripheral regions of Day 21 gemma cups (red rectangles) are shown at the bottom. (C). Schematic diagram showing the progression of gemma initiation in a gemma cup. The region where gemma initiation is occurring is colored green. (D). Localization of citrine fluorescence in cross-sections of Day 1, 7, 14, and 21 gemma cups in the *_pro_*Mp*KAI2B-CitNLS* line. Enlarged views of the periphery (yellow) and central (red) regions of the gemma cup floor are shown on the right side. Scale bars: 500 μm (B), 100 μm (D).

To test this hypothesis, we analyzed how gemma formation proceeds in a gemma cup in both WT and Mp*smxl* mutants (Figure 4B). On Day 1, when the gemma cup was first visible, gemmae initiation had already started in the central region of the gemma cup in both WT and Mp*smxl* mutants. On Day 4 and Day 7, gemma initiation continued in both WT and Mp*smxl*, gradually expanding the gemma initiation region to the peripheral area. No difference was observable between WT and Mp*smxl* until Day 7. However, on Day 14, no initiating gemma was observed in WT, while they were still observed in the peripheral region in the Mp*smxl* mutants. Gemma initiation at the peripheral region continued in the Day 21 gemma cup in Mp*smxl*. These observations revealed that gemma formation starts from the inner region of the gemma cup, and the gemma initiating area moves towards the peripheral region as the gemma cup grows. Then, gemma initiation stops at a certain time when the appropriate number of gemmae has been initiated in a gemma cup in WT (Figure 4C). We were interested to determine which of the gemma number in a cup or timing is a determinant for ending gemma initiation. To distinguish between these possibilities, we removed mature gemmae from the gemma cup every two days and analyzed the total number of gemmae formed. We assumed that if the number of gemmae in the cup is critical, their removal would lead to a delay in the end of gemma formation. We found that gemma formation stopped at around Day 10 even if mature gemmae were removed every other days (Figure S5A, B). Furthermore, the total number of gemmae formed in a cup did not differ significantly, whether gemma cups were removed or not (Figure S5C). This suggests that the time point, rather than the number of gemmae present in the gemma cup, is determinant in ending gemma formation.

We then analyzed the spatial localization of the activity of the KAI2-dependent signaling over 21 days using the _pro_*KAI2B*-CitNLS line as a marker (Figure 4D). Citrine fluorescence was observed in the whole region of the gemma cup floor cells and initiating gemmae on Day 1. On Day 7, all gemma cup floor cells and initiating gemmae showed fluorescence, while the fluorescence became weaker in the central region of the gemma cup. On Day 14, the fluorescence diminished in most areas of the floor of the gemma cup, with weak fluorescence in the peripheral region where gemma was sporadically initiating. Finally, on Day 21, the fluorescence disappeared in the whole region of the gemma cup floor. These observations indicate a good correlation between the localization of the KAI2-dependent signaling activity and gemma initiation.

In summary, we showed that gemma initiation starts from the inner region of the gemma cup and expands to the peripheral region of the floor of the gemma cup (Figure 4C). In WT, the KAI2-dependent signaling activity diminishes; thus, gemma initiation stops after a sufficient number of gemmae are initiated in the cup while it continues in Mp*smxl* loss of function mutant.

### *GEMMA CUP-ASSOCIATED MYB1* (*GCAM1*) works downstream of the KAI2-dependent signaling to activate gemma cup formation and gemma initiation

To identify genes involved in gemma initiation downstream of the KAI2-dependent signaling, we conducted RNAseq analysis and performed two comparisons. One was a comparison between Day 3 and Day 14 gemma cups of WT plants. Another comparison was between Day 14 WT and Day 14 Mp*smxl* gemma cups (Figure 5A; Data S1). More than 1,000 genes were found to be upregulated in both comparisons. Gene ontology (GO) enrichment analysis indicates that genes related to photosynthesis, light response, and chloroplast are enriched in the gemma cups actively initiating gemma compared to gemma cups that stopped gemma initiation (Figure S6). *GEMMA CUP-ASSOCIATED MYB1* (*GCAM1*), encoding an R2R3 MYB transcription factor, was identified among the commonly up-regulated DEGs (Data S1). Because GCAM1 is necessary for gemma cup initiation [6], we postulated that the KAI2-dependent signaling pathway controls gemma cup formation through the promotion of *GCAM1*. We first analyzed the genetic relationships between *GCAM1* and Mp*SMXL*. In the Mp*smxl gcam1* double mutants, gemma cup formation was inhibited entirely, supporting our view that *GCAM1* works downstream of Mp*SMXL* to control gemma cup initiation (Figure 5D).

**Figure 5.**
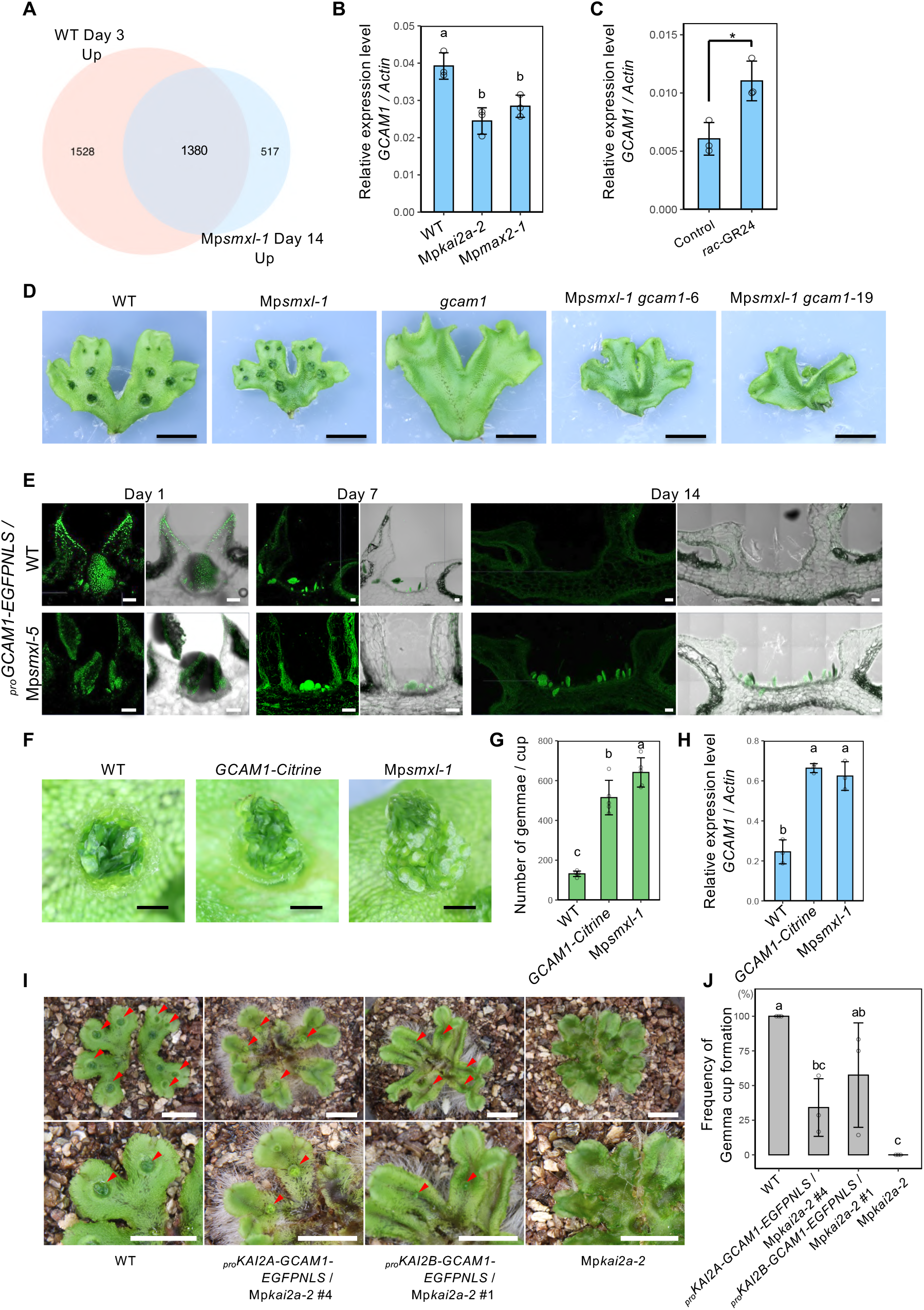
*GCAM1* works downstream of the KAI2-dependent signaling to activate gemma cup formation and gemma initiation. (A). Venn diagram showing the number of genes up-regulated (>1.5 x) in Day 3 WT gemma cups (pink) and Day 14 Mp*smxl-1* gemma cups (blue). (B). Expression level of Mp*GCAM1* relative to *M. polymorpha ACTIN* in the gemmae of one-day culture. Gemmae of WT, Mp*kai2a-2*, and Mp*max2-1* plants grown on 1/2 B5 medium for 24 hours were used for RNA extraction. Bars represent mean ±SD from three biological replicates. (C). Expression level of Mp*GCAM1* relative to *M. polymorpha ACTIN* in At*D14*^OX^/Mp*kai2a-4* plants treated with *rac*-GR24. Gemmae grown on 1/2 B5 medium for 24 hours, then transferred to 1/2 B5 liquid medium including Acetone (Control) or *rac*-GR24 (1 μM) and incubated for 6 hours were used for RNA extraction. Bars represent mean ±SD from three biological replicates. Significant differences between *rac-GR24* treated and untreated plants as determined by Student t-test with p < 0.05 are marked with an asterisk. (D). Three-week-old plants grown from the apical region of the thalli of WT, Mp*smxl*, Mp*gcam1*, and Mp*smxl* Mp*gcam1* double mutants. (E). Expression patterns of *_pro_*Mp*GCAM1-CitNLS* in Day 1, Day 7, and Day 14 gemma cups in WT (top) and Mp*smxl-1* (bottom) plants. (F). Gemmae in Day 14 gemma cups in WT, Mp*GCAM1-Citrine* (Yasui et al., 2019), and Mp*smxl-1*. (G). The number of gemmae per cup at Day 14 in WT, Mp*GCAM1-Citrine*, and Mp*smxl-1*. Bars represent the mean ±SD of five gemma cups. (H). The expression level of Mp*GCAM1* relative to *M. polymorpha ACTIN* in the gemmae of WT, Mp*GCAM1-Citrine*, and Mp*smxl-1*. Bars represent mean ±SD from three biological replicates. (I). Thalli of WT, Mp*kai2a-2*, and Mp*kai2a-2* expressing Mp*GCAM1* expression driven by Mp*KAI2A* or Mp*KAI2B* promoter (pro*KAI2A-GCAM1-EGFPNLS/kai2a-2* and pro*KAI2B-GCAM1-EGFPNLS/kai2a-2*, respectively) grown on soil. Red arrowheads indicate gemma cups. (J). Gemma cup formation frequency in WT, Mp*kai2a-2, proKAI2A-GCAM1-EGFPNLS/kai2a-2*, and pro*KAI2B-GCAM1-EGFPNLS/kai2a-2* grown on soil. The Tukey’s HSD test was used for multiple comparisons in (B), (G), (H), and (J). Statistical differences (p < 0.05) are indicated by different letters. Scale bars: 1 cm (D), 100 μm (D) and 1 mm (F) and (I).

To reveal the spatial and temporal expression pattern of *GCAM1*, we produced a marker line that expresses an *EGFP* fused with an NLS under the control of the *GCAM1* gene promoter (_pro_*GCAM1-EGFPNLS*). Fluorescence was observed in the initiating gemmae and the floor and frill of the gemma cup from Day 1 through Day 7 in WT (Figure 5E). This pattern resembled that of Mp*KAI2B*. Following the change of gemma initiation pattern over time in a gemma cup (Figure 4C), no fluorescence was observed in the frill of the gemma cup and the floor cells of the gemma cup on Day 14 in WT (Figure 5E). On the other hand, GCAM1 expression continued in the initiating gemma and the gemma cup floor cells on Day14 in Mp*smxl-1*, suggesting that Mp*SMXL* suppresses *GCAM1*.

Gemma cup formation does not occur in the *gcam1* loss-of-function mutants [6]. Therefore, it is not known whether *GCAM1* is involved in the control of gemma initiation or its growth. Reexamination of gemma and gemma cup formation in the *GCAM1-Citrine* knockin line previously reported revealed an increase in the gemma number to the same level as in Mp*smxl* in this particular line (Figures 5F, G). In accordance with the increase in the number of gemmae, *GCAM1* expression was enhanced in the *GCAM1-Citrine* knockin line (Figure 5H). This indicates that an increase in *GCAM1* expression causes an increase in the gemma number and suggests that *GCAM1* is involved in the control of gemma initiation in addition to gemma cup formation. Moreover, the expression of *GCAM1* was enhanced in the Mp*smxl* mutants (Figure 5H). This supports the view that *GCAM1* is a downstream regulator of the KAI2-dependent signaling pathway (Figure 5H).

Finally, we tested whether the *GCAM1* rescues the defects in gemma cup formation in the Mp*kai2a* mutants. *GCAM1* fused with the EGFP gene under the control of either Mp*KAI2A* or Mp*KAI2B* promoters was introduced in the Mp*kai2a* mutant (Figures 5I). When grown on soil, gemma cup formation was suppressed in Mp*kai2a-2* (Figures 1C, 5I). The expression of *GCAM1* rescued this defect (Figures 5I, J). These results indicate that *GCAM1* is sufficient to rescue gemma cup formation in Mp*kai2a-2*. On the other hand, other defects observed in the signaling mutants, namely upward growth of thallus and weakened dormancy in the dark, were not complemented by the expression of *GCAM1* (Figure S7). These results support our hypothesis that *GCAM1* is a downstream regulator of the KAI2-dependent signaling pathway, which is specific to the control of gemma cup formation and gemma initiation.

### KAI2-dependent signaling is not involved in the control of axillary meristem formation in rice and *Arabidopsis*

We first tested if the control of gemma initiation by KAI2-dependent signaling is conserved in *M. paleacea*, a species closely related to *M. polymorpha*. An increased number of gemmae were observed in the mutants of the *SMXL* gene in *M. paleacea*, indicating that the control of gemma formation by the KAI2-dependent signaling is conserved (Figure 6A).

**Figure 6.**
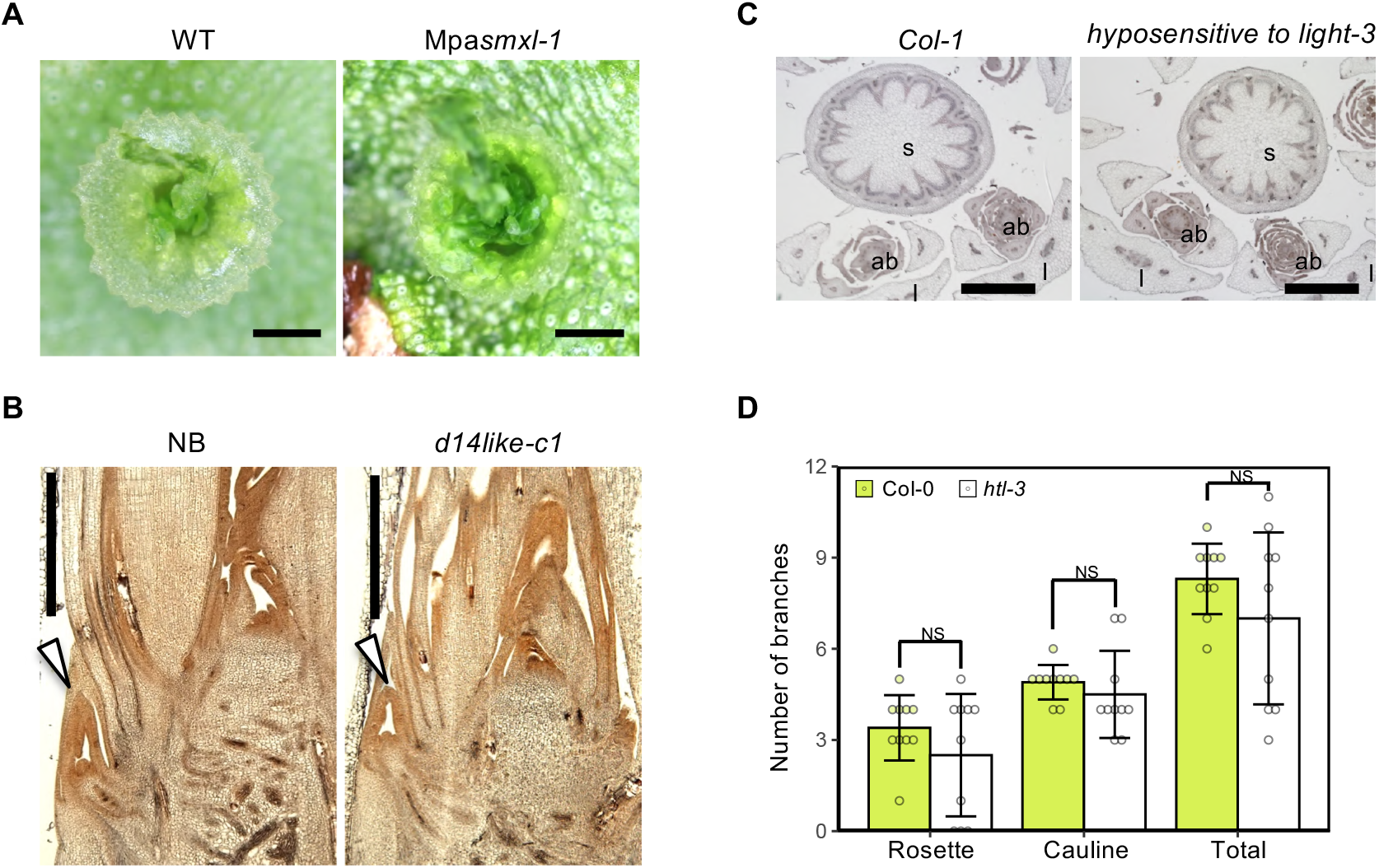
KAI2-dependent signaling is not involved in the control of axillary meristem formation in rice and Arabidopsis. (A). Mpa*smxl* mutant in *Marchantia paleacea*. The number of gemmae in a cup is increased in Mpa*smxl*. More than five plants were analyzed, and all showed a similar phenotype. (B). Phenotype of rice *d14like-c1* (*d14l-c1*) mutant. Axillary bud (arrowheads) formation is not affected in *d14l-c1*. More than six plants were analyzed, and all showed a similar phenotype. (C). Phenotype of *Arabidopsis hyposensitive to light* (*htl*) mutant. Axillary bud formation is not affected in *htl-3*. s, stem; ab, axillary bud; l, leaf. Five plants were sectioned and analyzed, and all showed a similar phenotype. (D). Number of branches in WT and in the *htl* mutant. Branches formed in the axil of cauline leaves and rosette leaves six weeks after germination were counted. Bars represent the mean ± SD of ten Arabidopsis plants. NS indicates no significant difference in the Student’s t-test at p < 0.05. Scale bars: 1 mm (A) to (C).

Close homologs of *GCAM1* are required for suppression of axillary bud initiation in angiosperms, including tomato and *Arabidopsis* [8–10]. Therefore, we anticipated that if the genetic module consisting of the KAI2-dependent signaling pathway and GCAM1 is conserved in seed plants, axillary meristem formation will be affected in KL signaling mutants. However, axillary bud formation was not affected in *hyposensitive to light-3* or in *d14like-c1*, KAI2 mutants in *Arabidopsis* and rice, respectively [38, 45] (Figures 6B to D).

## Discussion

### The number of gemmae in a gemma cup is controlled through ON/OFF switching of KAI2-dependent signaling

Controlling the number of offspring is crucial for successful reproduction for all creatures. Thus, all species developed species-specific reproductive strategies. The finding of this study that the defined number of gemmae are formed in a gemma cup is a significant first step toward understanding the principle of the reproductive strategy of *M. polymorpha*. We revealed that the KAI2-dependent signaling promotes gemma cup formation and gemma initiation. Moreover, the role of this signaling is not a simple promotion of these two steps. Intriguingly, the number of gemmae in a cup is controlled through ON/OFF switching of the KAI2-dependent signaling. We showed that gemma formation starts from the central part of the gemma cup and proceeds to the peripheral region as the gemma cup grows. Then, gemma initiation stops at a certain time; thus, the number of gemmae formed in the gemma cup is controlled by halting the signaling at a particular time when a proper number of gemmae has been initiated. This finding raised many questions that warrant further study. First of all, what determines the end of gemma formation needs to be clarified. There are two possibilities: the number of gemmae formed and the developmental timing. We showed that the removal of gemmae from the gemma cup did not affect the timing of the termination of new gemma initiation. The total number of gemmae formed was similar to that of the gemma cups without the removal treatment. Thus the number of gemmae present in a cup is unlikely to be determinant. However, we cannot rule out the possibility that the total number of gemmae initiated in the gemma cup is critical for the timing of the end of gemma formation. If plants stop new gemma initiation based on the total number of gemmae initiated, mechanisms that enable plants to sense and remember the number of gemmae formed would be needed. Identifying the nature of the mechanisms that control the developmental signals is the next challenge. Whichever hypothesis is correct, these mechanisms are totally unknown.

### GCAM1 works downstream of the KAI2-dependent signaling to control vegetative reproduction in *M. polymorpha*

We previously reported that MpKAI2A acts as a receptor of a yet uncharacterized ligand in the KAI2-dependent signaling pathway of *M. polymorpha* and transduces the signal through the degradation of MpSMXL, which works as a suppressor of downstream gene regulatory cascades [43]. SMAX1/SMXL genes, members of a small gene family, function as downstream suppressors of the D14-dependent strigolactone signaling pathway and KAI2-dependent KAR/KL signaling pathway [23–26, 28]. Our knowledge about genes downstream of SMAX1/SMXL is still limited, even in angiosperm species [46–48]. This study revealed that GCAM1 works in the downstream genetic cascade of the KAI2-dependent signaling pathway in *M. polymorpha*. It is not known whether MpSMXL directly suppresses GCAM1, but the attenuation of the KAI2-dependent signaling pathway allows the accumulation of MpSMXL, leading to the suppression of GCAM1.

*GCAM1* was reported as a regulator of gemma cup formation in *M. polymorpha* [6]. However, it was not known whether *GCAM1* is also involved in gemma formation due to the severe phenotypes of *gcam1* loss of function mutants in which gemma cup formation, the step preceding gemma initiation, is wholly suppressed. Here, we showed that a slight increase in *GCAM1* expression caused an increase in gemma number, indicating that *GCAM1* activates gemma initiation in addition to gemma cup formation. Thus, *GCAM1* likely regulates a biological process involved in these two steps of gemma development. A gemma cup originates from a few cells derived from the dorsal and lateral merophytes, and is potentially formed at each point of bifurcation of the thallus tip [4].

In contrast, a gemma initiates from a cell on the floor of the gemma cup [4, 5]. Cells specified as gemma initials undergo cell divisions to form gemma while other cells at the bottom of the gemma cup are maintained as the floor cells or specified as mucilage cells. Overexpression of *GCAM1* by the EF1 promoter caused an excessive propagation of undifferentiated cell masses [6]. Based on this observation, it was proposed that the function of *GCAM1* may be to specify the undifferentiated state of cells. Therefore, we hypothesize that the cells need to be undifferentiated before they can be specified as gemma initial cells and gemma cup founder cells, and *GCAM1* is required for these processes. Understanding the undifferentiated state of cells conferred by GCAM1 at the molecular level will be crucial to elucidate the molecular mechanisms underlying the vegetative reproduction of *M. polymorpha*. Precise determination of the spatial and temporal requirements of *GCAM1* expression for gemma cup formation and gemma initiation will be critical.

We showed that *GCAM1* controls both gemma cup formation and gemma formation. On the other hand, Mp*CLE2* regulates only gemma cup formation but not gemma formation, *KAR* is required only for gemma initiation, and *RSL1* is required for specifying gemma initial and mucilage cells but not for gemma cup formation [11, 13]. Gemma cup formation and gemma initiation are complex processes regulated by numerous genes and gene modules that are either specific to each process or common to both processes.

### Evolution of KAI2-dependent signaling as a regulator of vegetative reproduction

We showed that *GCAM1* works downstream of the KAI2-dependent signaling in *M. polymorpha*. In flowering plants, KAI2-dependent signaling controls variable traits, including seed germination, hypocotyl development, leaf development, root hair and root development, and drought tolerance. However, to our knowledge, the involvement of the KAI2-dependent signaling in the control of axillary bud formation or vegetative reproduction has not been reported. Indeed, we showed that axillary bud formation is not affected in the loss of function mutants of KAI2 in rice and *Arabidopsis*. Therefore, despite the conservation of GCAM1 orthologs’ function in the control of vegetative reproduction (6, 8-10), the link between GCAM1 and the KAI2-dependent signaling was lost in the lineage leading to seed plants or acquired in the lineage leading to *M. polymorpha*. Analysis of other bryophytes, such as mosses and hornworts, and distantly diverging lineages of extant vascular plants will be needed to establish which of these events took place.

In angiosperms, the shoot branching pattern is mainly controlled via axillary bud outgrowth rather than axillary bud formation [49, 50]. Axillary bud formation is largely developmentally defined rather than being environmentally responsive. SL was first identified as a plant hormone that inhibits axillary bud outgrowth in angiosperms [51, 52]. Later, it was demonstrated that the SL signaling pathway adjusts various aspects of growth and development to environmental conditions. The function of SLs as a class of phytohormones was acquired in the common ancestor of seed plants through the evolution of D14, the cognate receptor, by duplication of *KAI2* [40, 52]. Thus, control of shoot branching by the hormonal function of SLs occurs only in seed plants. On the other hand, shoot branching through axillary meristem formation is a widespread phenomenon observed in land plants. One of the advantages of controlling shoot branching at the level of bud outgrowth is that it enables rapid responses to changes in environmental conditions. An intriguing possibility to be tested in the future is that SL signaling replaced KAI2-dependent signaling as a regulator of shoot branching.

### Environmental control of vegetative reproduction in *M. polymorpha*

Environmental conditions influence the growth and development of plants. Nutrient availability is one of the significant environmental factors affecting plant growth and development. We showed that among the three nutrient deficiencies analyzed, K deficiency is the one that affects gemma cup formation in *M. polymorpha*. Our analysis indicates that the potassium-dependent control of gemma cup formation is independent of the KAI2 pathway (Figure 7). Potassium is a crucial modulator of cell homeostasis and is required for many biological processes, such as the activation of more than 60 enzymes, tolerance to abiotic stresses, and stomata regulation. Low K+ concentration controls root and shoot development responses through ethylene signal transduction [53]. In addition, the link between K and other hormones, including brassinosteroids, auxins, gibberellins, jasmonic acid, and cytokinins, was reported in the control of abiotic stress tolerance. However, the involvement of K in the control of vegetative reproduction has not been demonstrated. Understanding how the information of K availability is transduced into the process of gemma cup formation will be crucial for our understanding of the control of vegetative reproduction in *M. polymorpha*.

**Figure 7.**
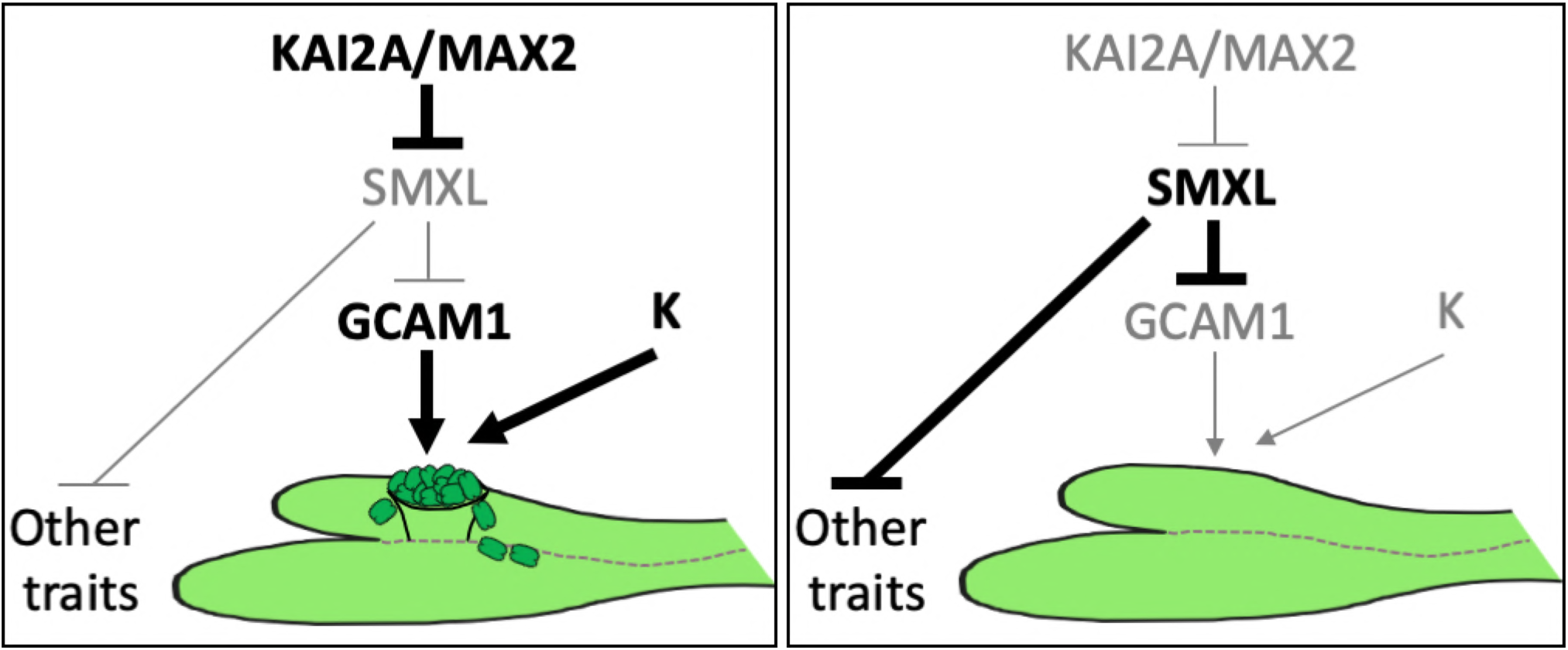
Model of the control of vegetative reproduction of *M. polymorpha* by the KAI2-dependent signaling. MpKAI2A, MpMAX2, and MpSMXL work in a single signaling pathway, which depends on the degradation of MpSMXL. This signaling pathway positively controls gemma cup formation and gemma initiation. The number of gemmae in a cup is controlled through the ON/OFF switching of this signaling. *GCAM1* is a downstream regulator of the KAI2-dependent signaling pathway in controlling gemma cup formation and gemma initiation. K also positively controls gemma cup formation independently of the KAI2-dependent signaling pathway.

KAI2-mediated signaling regulates low P-dependent root and root hair development in *Arabidopsis thaliana* and *Lotus japonica* [34, 36, 54]. In both species, ethylene biosynthesis and signaling are regulated by the KAI2-dependent signaling, leading to changes in auxin accumulation. In the control of root and root hair development, the KAI2 pathway promotes the Low-P responses. In contrast to these studies, we did not find a significant link between the availability of P or N and gemma cup formation. Our findings indicate that *M. polymorpha* plants modulate the KAI2-dependent signaling, leading to the formation of an adequate number of gemma cups and gemmae.

Previously, we showed that the KAI2 pathway positively controls thallus growth, planar growth of the thallus, and dormancy in the dark. Horizontal growth of large thalli on the ground is advantageous for efficient photosynthesis, and keeping them in the dormant state in darkness is essential to preserve energy. In this study, we established modulation of vegetative reproduction as a role of the KAI2-dependent signaling. Overall, it is likely that KAI2-dependent signaling regulates growth and development in response to environmental cues in *M. polymorpha* to optimize proliferation. The next challenge is to understand the environmental factors that are is integrated into the KAI2-dependent signaling to optimize vegetative reproduction in *M. polymorpha*.

