## Supplemental Figures for "*KARRIKIN INSENSITIVE2* (*KAI2*)-dependent signaling pathway controls vegetative reproduction in *Marchantia polymorpha*"

Figure S1.

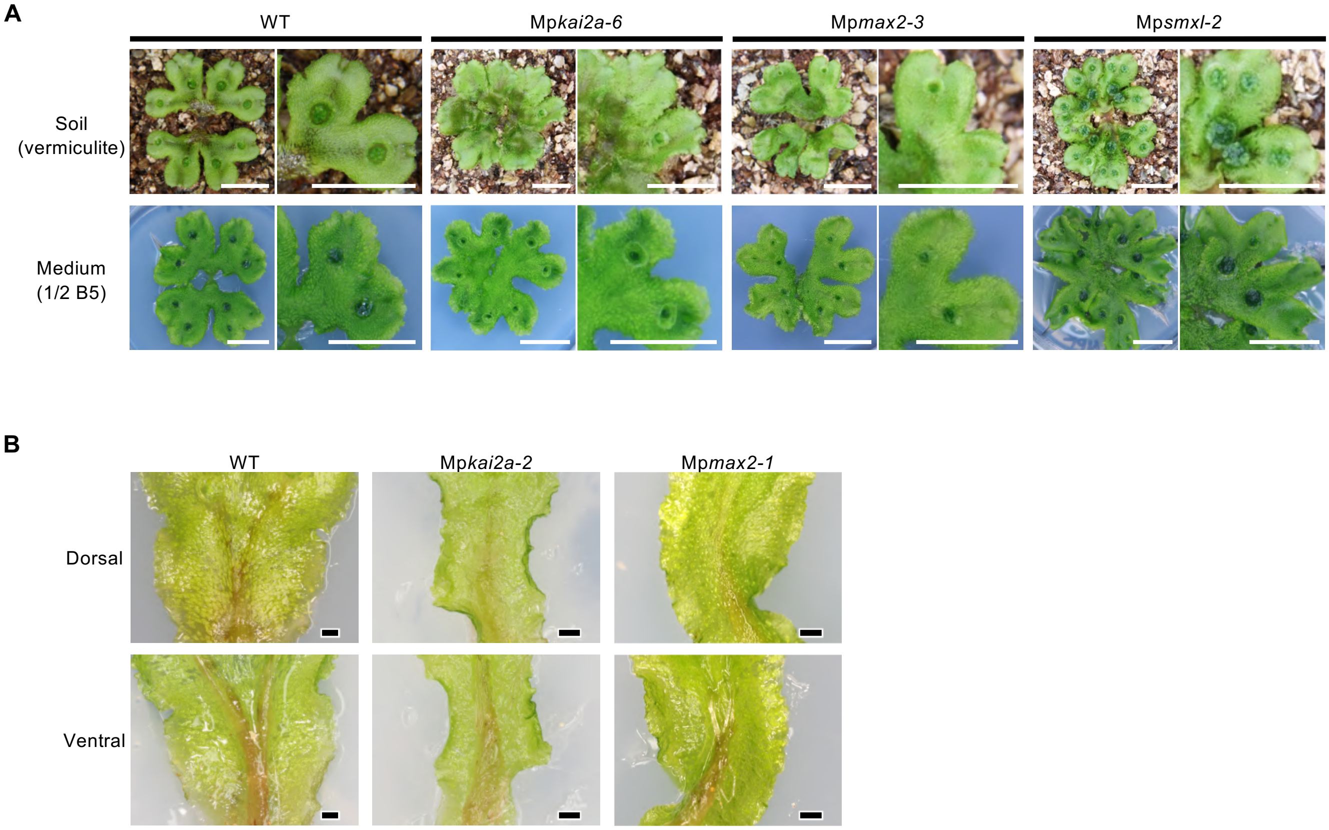


**Figure S1. Phenotypes of KAI2-dependent signaling mutants.**
(A). Gemma cup phenotypes of WT, Mp*kai2a-6,* Mp*max2-3*, and Mp*smxl-2* grown on soil or synthetic medium. Gemmae were grown on 1/2 B5 medium for six days, then transferred to soil (upper panel) or a fresh 1/2 B5 medium (lower panel) and grown for another 14 days before taking the images.
(B). Midrib phenotypes of WT, Mp*kai2a-2*, and Mp*max2-1*. WT, Mp*kai2a-2*, and Mp*max2-1* gemmae were grown on 1/2 B5 medium for six days, then transferred to soil and grown for another 6 weeks.
Scale bars: 1 cm (A) and 1 mm (B).

Figure S2.

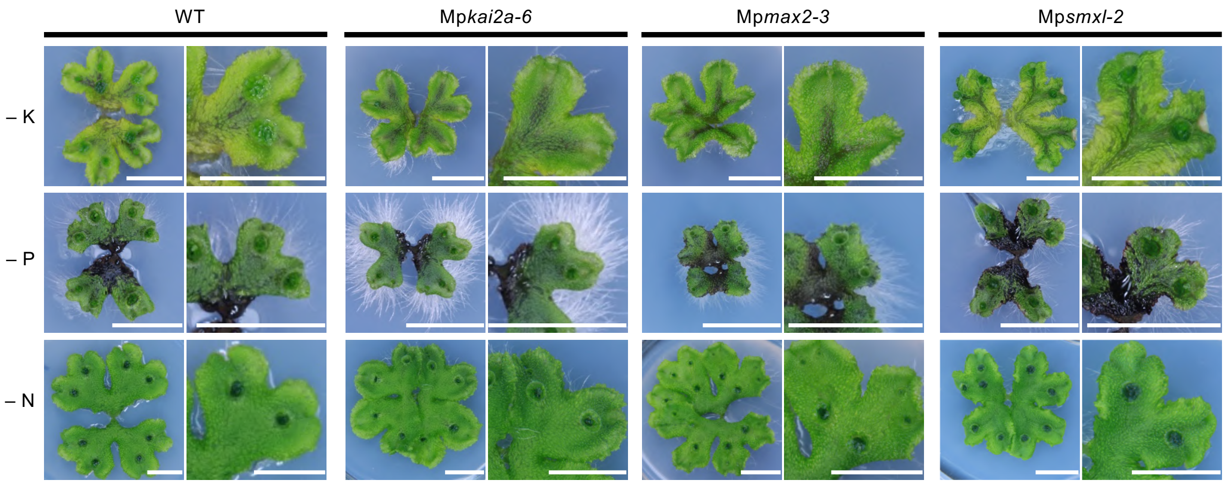


**Figure S2. Effect of nutrient deficiency and mutations on the KAI2-dependent signaling pathway to gemma cup formation.**
WT, Mp*kai2a-6,* Mp*max2-3*, and Mp*smxl-2* plants were grown from a gemma on 1/2 B5 medium for six days, then transferred to a medium lacking potassium (-K), phosphate (-P), or nitrogen (-N) and grown for another 14 days. A close-up view of the thallus is shown on the right side. Scale bars: 1 cm.

Figure S3.

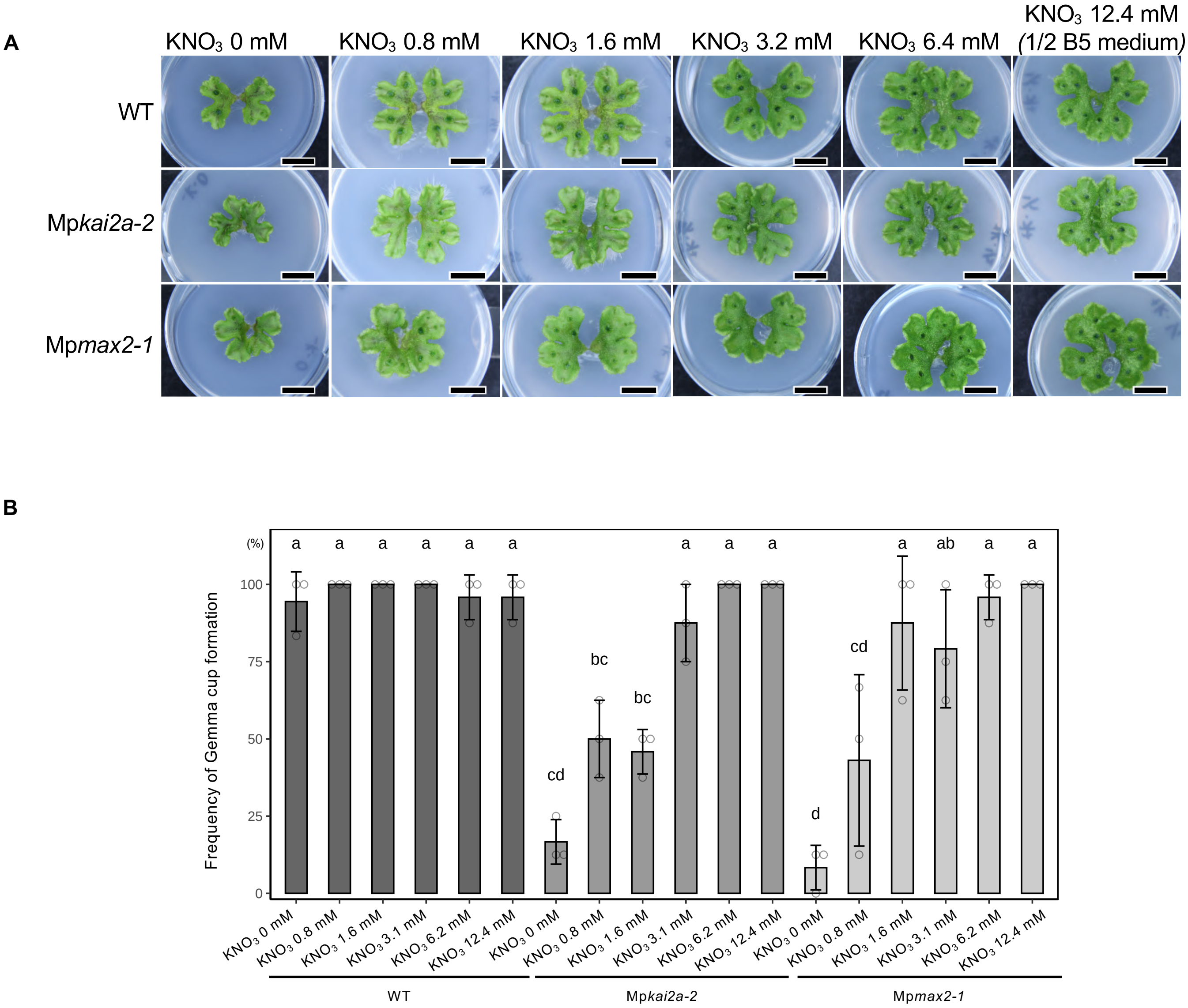


**Figure S3. Effect of potassium (-K) deficiency on gemma cup formation.**

(A). Gemma cup phenotypes of KAI2-dependent signaling mutants grown on synthetic medium. WT, Mp*kai2a-2*, or Mp*max2-1*, gemmae were grown on 1/2 B5 medium for six days. Plants were then transferred to 1/2 B5 medium containing different concentrations of potassium (0 to 12.4 mM $KNO_3$) and grown for another 14 - 21 days. Scale bars: 1 cm.

(B). Frequency of actual gemma cup formation per eight possible gemma cup formation points after 3rd bifurcation in WT (green), Mp*kai2a-2*(grey), and Mp*max2-1*(white) grown on 1/2 B5 medium with different concentrations of $KNO_3$. Bars represent mean ± SD (n = 3). Tukey's HSD test was used for multiple comparisons. Statistical differences ($p < 0.05$) are indicated by different letters.

Figure S4.

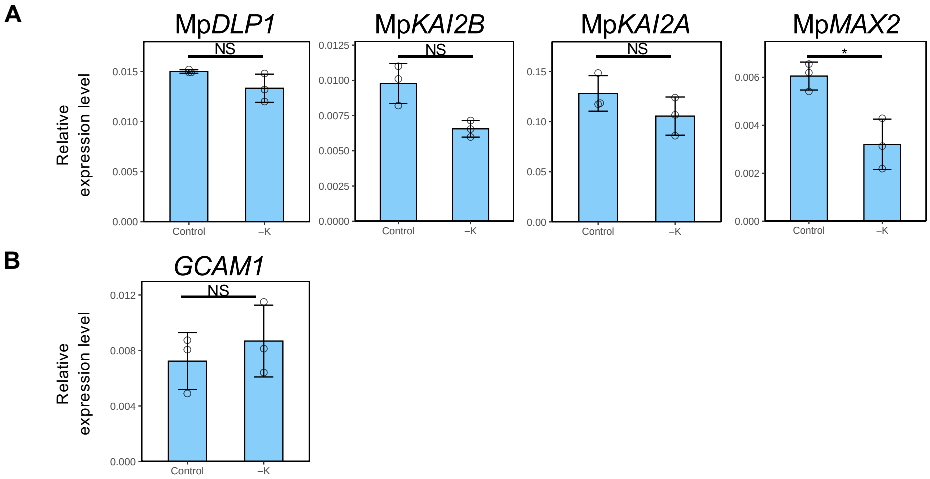


**Figure S4. Effects of potassium (-K) deficiency on the KAI2-dependent signaling pathway.**
**(**A)-(B), WT plants were grown on 1/2 B5 medium for seven days, then transferred to 1/2 B5 liquid medium lacking potassium (-K) and cultured for 24 hours. The expression levels of Mp*DLP1,* Mp*KAI2B,* Mp*KAI2A*, Mp*MAX2* (A)*,* and *GCAM1* (B) relative to the Mp*Actin* gene as determined by q-PCR are shown.
Bars represent mean ±SD from three biological replicates. Asterisks indicate significant differences relative to the control (*$p < 0.05$, NS = no significant difference).

Figure S5.

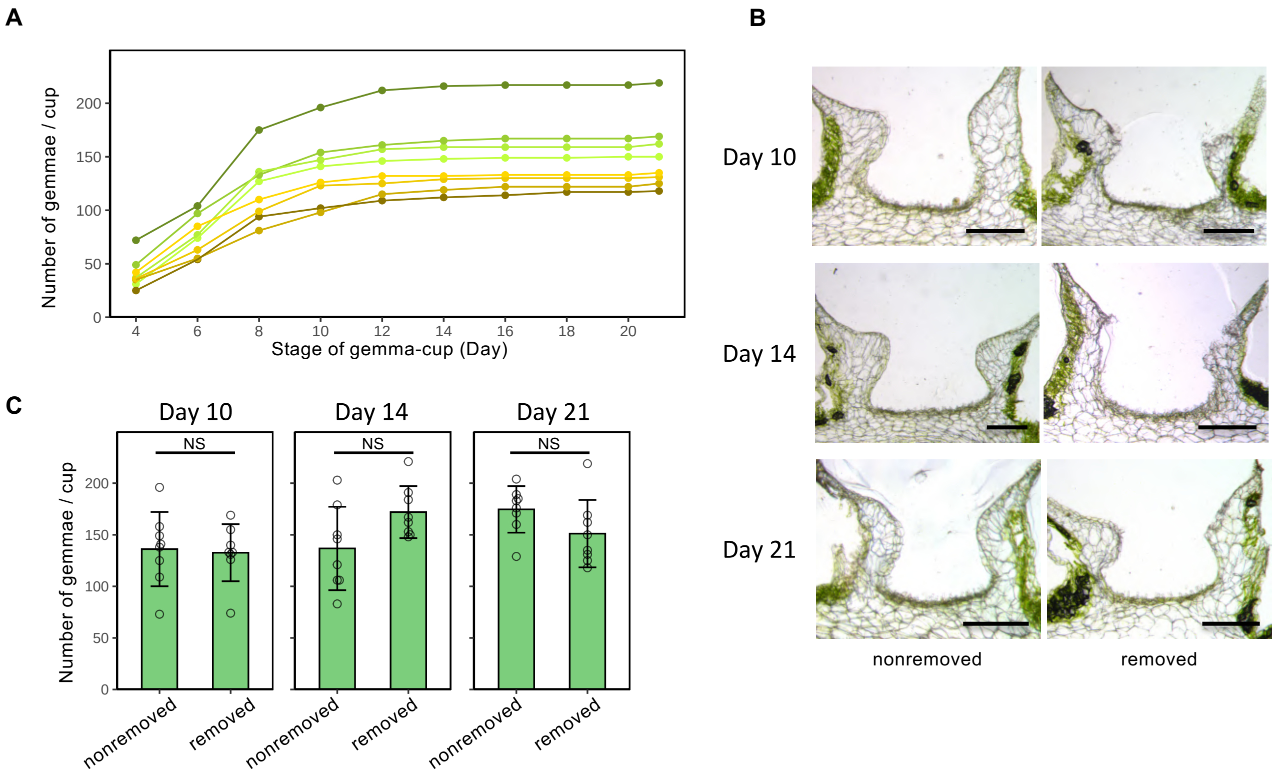


**Figure S5. Effect of gemma removal on gemma formation in a gemma cup.**
(A). Time-course of gemma formation in gemma-cups of WT from which mature gemmae were removed. The first day when the gemma cup becomes visible is designated as Day 1. The gemmae were removed every two days from Day 4 to Day 20. Each colored line indicates the result of an independent gemma cup.
(B). Cross-sections of the gemma cups of WT with and without gemma removal on Day 10, 14, and 21. Scale bars: 500 μm.
(C). The total number of gemmae formed in a gemma cup of WT with (white) and without (green) gemma removal. Bars represent mean ±SD from eight gemma cups of Day 10, 14, or 21. NS indicates no significant difference in the Student's t-test at $p < 0.05$.

Figure S6.

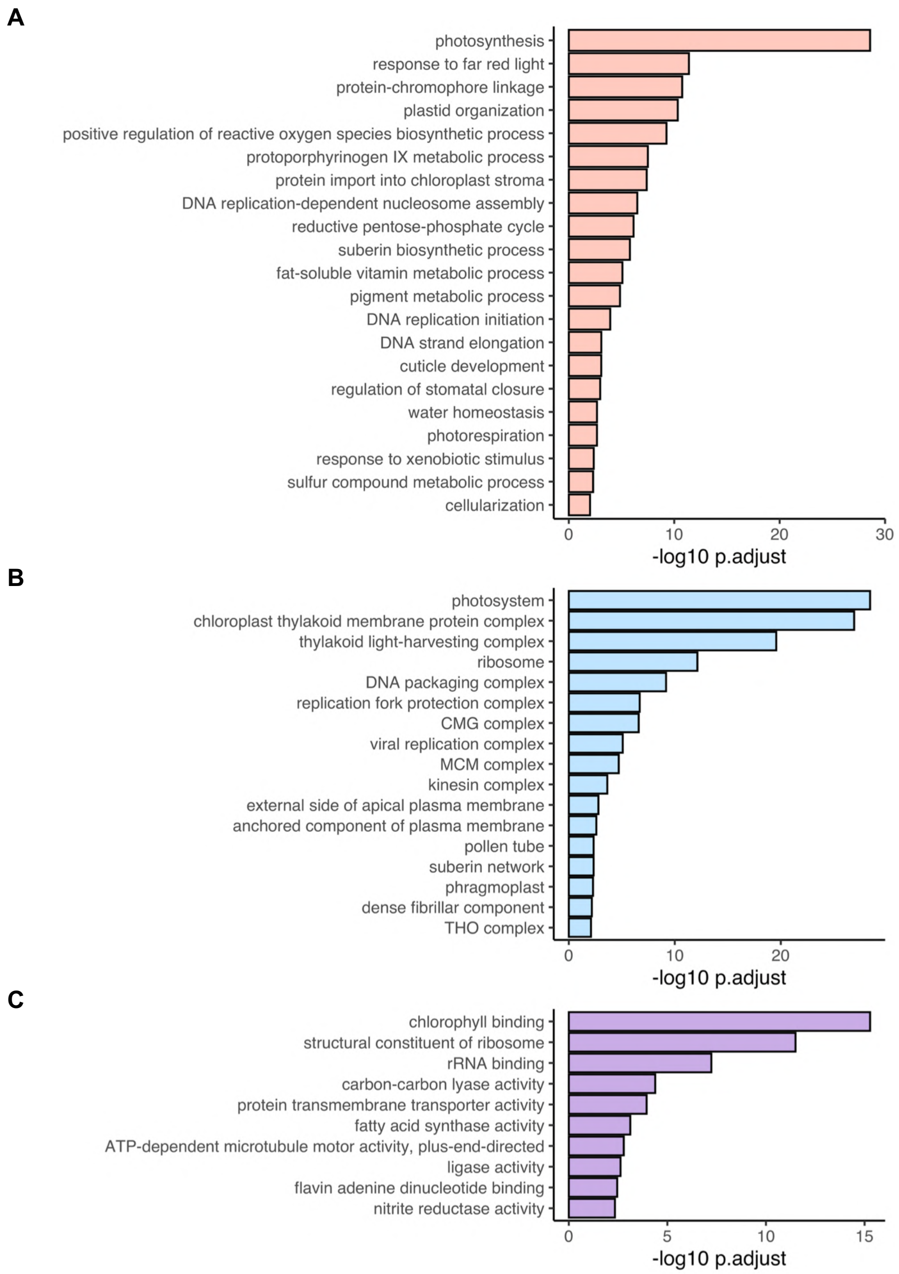


**Figure S6. Gene ontology (GO) enrichment analysis of up-regulated genes in gemma initiating gemma-cups, as shown in Figure 5A.**
(A). GO Biological Process genes
(B). GO Cellular Component genes
(C). GO Molecular Function genes
The y-axis represents annotations of GO terms. The x-axis represents the minus logarithm of the p-value. GO term annotation and enrichment analysis were performed according to the methods described previously (Ishida et al., 2022).

Figure S7.

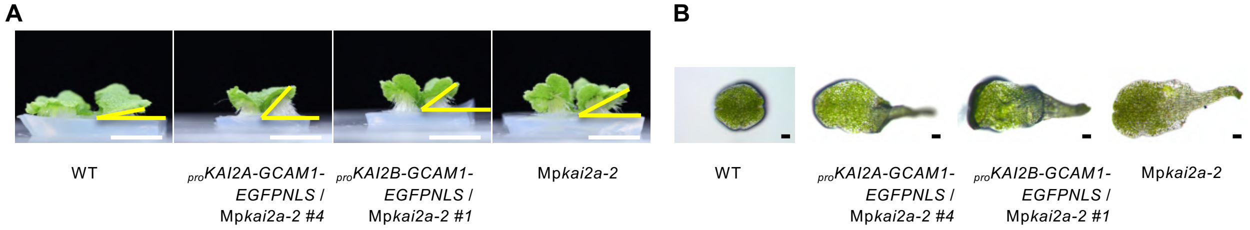


**Figure S7. WT, Mp*kai2a-2,* and Mp*kai2a-2* plants with Mp*GCAM1* expression driven by the Mp*KAI2A* or Mp*KAI2B* promoters.**
(A). Side view of plants. The upward-growth phenotype of the Mp*kai2a* mutant was not rescued by the expression of Mp*GCAM1*. The plants were grown from gemmae on 1/2 B5 medium for 14 days.
(B). The release-of-dormancy phenotype of the Mp*kai2a* mutant was not rescued by the expression of Mp*GCAM1* driven by the Mp*KAI2A* or Mp*KAI2B* promoters. The plants were grown from gemmae in the dark for 14 days.
Scale bars: 1 cm (A), 100 μm (B).

**Supplemental Table 1.** Production of CRISPR mutants.

| Allele | Gene | gRNA sequence | Mutatuin site | Amino acid |
|---|---|---|---|---|
| Mp*kai2a-6* | Mp*KAI2A* | GGAGATCAGCTCGTGGTACT | g.59_60insTTAGGACAGT CAGCTCGTCAGC | p.L21X |
| Mp*max2-3* | Mp*MAX2* | GGGACTGGTACCTCTTGGAG | g.–729_129delins TCTTCCTC | No start codon |
| At*D14*$^{ox}$/Mp*kai2a-5* | Mp*KAI2A* | TGTACGATACCATGGGAGCA | g.155_156insT | p.A53fsX62 |
| Mp*smxl-1 gcam1-6* | Mp*GCAM1* | GTCCTTGGTCGCCCGAAG | g.123_507del | p.K42fsX441 |
| Mp*smxl-1 gcam1-19* | Mp*GCAM1* | GTCCTTGGTCGCCCGAAG | g.–543_143 delins GAGGATGCA | No start codon |
| *proGCAM1-EGFPNLS /* Mp*smxl-5* | Mp*SMXL* | GTGTCCGAGGCAAAGAGTCG | g.81_93del | p.K28fsX36 |
| Mpa*smxl*-1 | Mpa*SMXL* | TTCCGAGGCAAAGAGTCG | g.86_87insA | p.S29fsX33 |

※ Notations were followed den Dunnen JT and Antonarakis SE (2000). Hum.mutat. 15:7-12

**Supplemental Table 2.** Primers used in this study.

| Name | Sequence (5'→3') | Usage |
|---|---|---|
| MpACT-qPCR-F | AGGCATCTGGTATCCACGAG | qRT-PCR control |
| MpACT-qPCR-R | ACATGGTCGTTCCTCCAGAC | qRT-PCR control |
| GCAM1-qPCR-F | GATGTCGGGACGCAAATAAT | qRT-PCR |
| GCAM1-qPCR-R | TGTCATTGCGTAGGGAGATTC | qRT-PCR |
| Mp*PHT1*-qPCR-F | TGCGGTGGTAACCATTTAGC | qRT-PCR |
| Mp*PHT1*-qPCR-R | GCACGATGAATGTTGTGGAG | qRT-PCR |
| Mp*NRT2.1*-qPCR-F | CACCCTGCCAATTATCTTCG | qRT-PCR |
| Mp*NRT2.1*-qPCR-R | CATTGGCTTGCTCTTCCTTG | qRT-PCR |
| Mp*DLP1*-qPCR-F | GGTGTGAAGAAAGTTGGAGTTATGG | qRT-PCR |
| Mp*DLP1*-qPCR-R | GTGTGAGGAATGAGGGATGGTT | qRT-PCR |
| Mp*KAI2A*-qPCR-F | GTATGATCGGGTGCCTTGC | qRT-PCR |
| Mp*KAI2A*-qPCR-R | TGCATTGCCTCAAAGAGCTG | qRT-PCR |
| Mp*MAX2*-qPCR-F | GTATCCCCGATTGAATTTCG | qRT-PCR |
| Mp*MAX2*-qPCR-R | GATTCCCGCAACACAAGAG | qRT-PCR |
| proKAI2A-F | caccGCCACGCCAGACTAGAGATC | Construction of $_{pro}$*KAI2A-CitrineNLS* |
| proKAI2A-R | ACCACCGCTTTGCTATGGTT | Construction of $_{pro}$*KAI2A-CitrineNLS* |
| proKAI2B-F | caccTGTGAAGAAGCTTCCCGTGG | Construction of $_{pro}$*KAI2B-CitrineNLS* |
| proKAI2B-R | TGTAGCTCTGGATGCAGTCAG | Construction of $_{pro}$*KAI2B-CitrineNLS* |
| MpGCAM1-gRNA1-F | ctcgGTCCTTGGTCGCCCGAAG | Construction of the genome editing vector for *GCAM1* |
| MpGCAM1-gRNA1-R | *aaacCTTCGGGCGACCAAGGAC* | *Construction of the genome editing vector for GCAM1* |
| CRISPR-check_L | CCTGTAATGAGTGGATTCGCTTG | For checking of the edited site for *GCAM1* |
| CRISPR-check_R | TAGGACTCGATGCTGAACTCGTC | For checking of the edited site for *GCAM1* |
| proGCAM1_L0 | caccCGATCACCTCTCGTCCCATG | Construction of $_{pro}$*GCAM1-EGFPNLS* |
| prpGCAM1_R1 | GAATGTGGATCACGGTCGGG | Construction of $_{pro}$*GCAM1-EGFPNLS* |
| MpD53CRISPR270for | gcacccagcctctcgGGACAGATTGCCATCGAACCg ttttagagctagaa | Construction of the genome editing vector for Mp*SMXL* |
| MpD53CRISPR270rev | ttctagctctaaaacGGTTCGATGGCAATCTGTCCcg agaggctgggtgc | Construction of the genome editing vector for Mp*SMXL* |
| MpD53 CRISPR90For | gcacccagcctctcgGTGTCCGAGGCAAAGAGTCG gttttagagctagaa | Construction of the genome editing vector for Mp*SMXL* |
| MpD53CRISPR90Rev | ttctagctctaaaacCGACTCTTTGCCTCGGACACcg agaggctgggtgc | Construction of the genome editing vector for Mp*SMXL* |
| MpD53genotypefor | TTGGACGCCCGGTCGAGCTTTTC | For checking of the edited site for Mp*SMXL* |
| MpD53genotype2rev | GAAAGATGGCGAAAGACTAGA | For checking of the edited site for Mp*SMXL* |
| MpaSMXL_CR1_F | CTCGTTCCGAGGCAAAGAGTCG | Construction of the genome editing vector for Mpa*SMXL* in *Marchantia paleasea* |
| MpaSMXL_CR1_R | AAACCGACTCTTTGCCTCGGAA | Construction of the genome editing vector for Mpa*SMXL* in *Marchantia paleasea* |
| MpaSMXL_geno_F | TCCAGCACACTTTAACCCCG | For checking of the edited site for Mpa*SMXL* |
| MpaSMXL_geno_R | GTCAGAGATGGCGAGGACTG | For checking of the edited site for Mpa*SMXL* |
